# The genetic legacy of Zoroastrianism in Iran and India: Insights into population structure, gene flow and selection

**DOI:** 10.1101/128272

**Authors:** Saioa López, Mark G. Thomas, Lucy van Dorp, Naser Ansari-Pour, Sarah Stewart, Abigail L. Jones, Erik Jelinek, Lounès Chikhi, Tudor Parfitt, Neil Bradman, Michael E. Weale, Garrett Hellenthal

## Abstract

Zoroastrianism is one of the oldest extant religions in the world, originating in Persia (present-day Iran) during the second millennium BCE. Historical records indicate that migrants from Persia brought Zoroastrianism to India, but there is debate over the timing of these migrations. Here we present novel genome-wide autosomal, Y-chromosome and mitochondrial data from Iranian and Indian Zoroastrians and neighbouring modern-day Indian and Iranian populations to conduct the first genome-wide genetic analysis in these groups. Using powerful haplotype-based techniques, we show that Zoroastrians in Iran and India show increased genetic homogeneity relative to other sampled groups in their respective countries, consistent with their current practices of endogamy. Despite this, we show that Indian Zoroastrians (Parsis) intermixed with local groups sometime after their arrival in India, dating this mixture to 690-1390 CE and providing strong evidence that the migrating group was largely comprised of Zoroastrian males. By exploiting the rich information in DNA from ancient human remains, we also highlight admixture in the ancestors of Iranian Zoroastrians dated to 570 BCE-746 CE, older than admixture seen in any other sampled Iranian group, consistent with a long-standing isolation of Zoroastrians from outside groups. Finally, we report genomic regions showing signatures of positive selection in present-day Zoroastrians that might correlate to the prevalence of particular diseases amongst these communities.

## Introduction

The Zoroastrian religion developed from an ancient religion that was once shared by the ancestors of tribes that settled in Iran and northern India. It is thought to have been founded by the prophet priest Zarathushtra (Greek, Zoroaster). Most scholars now believe he lived around 1200 BCE, at a time when the ancient Iranians inhabited the areas of the Inner Asian Steppes prior to the great migrations south to modern Iran, Afghanistan, Northern Iraq and parts of Central Asia. Zoroastrianism became the state religion of three great Iranian empires: Achaemenid (559-330 BCE) founded by King Cyrus the Great and ended by the conquest of Alexander the Great, Parthian (c. 247 BCE - 224 CE), and Sasanian (224-651 CE), during which time the religion as an imperial faith is best known. Zoroastrianism ceased to be the state religion of Iran after the Arab conquests (636-652 CE), although it is thought that widespread conversion to Islam did not begin until about 767 CE^1^.

According to Parsi (i.e. Indian Zoroastrians) tradition, a group of Zoroastrians set sail from Iran to escape persecution by the Muslim majority. They landed on the coast of Gujarat (India) where they were permitted to stay and practice their religion. The date of the arrival remains has been the cause of speculation and varies between 785 CE^2^ and 936 CE^3^. These dates, among others, are based on the *Qisseh-ye Sanjan,* a legendary account of the journey by sea from Iran and settlement in India^4^. However, maritime trade is known to have taken place between ethnic groups from Iran, including Zoroastrians, and peoples in India long before the arrival of Islam^5^. Down the subsequent centuries, the Indian Zoroastrians (also known as Parsis) maintained contact with the Zoroastrians of Iran and later became an influential minority under British Colonial rule.

Zoroastrian communities today are concentrated in India (61,000), Southern Pakistan (1,675) and Iran - mainly in Tehran, Yazd and Kerman – (14,000). In the last 200 years Zoroastrians, both Parsi and Irani, have formed diaspora communities in North America (14,306), Canada (6,422), Britain (5,000), Australasia (3,808) and the Middle East (2,030). Zoroastrianism is a non-proselytising religion, with a hereditary male priesthood of uncertain origins^6^. Among the Parsis, priestly families are distinguished from the laity. Priestly status is patrilineal, although there is also a strong matrilineal component with the daughters of priests encouraged to marry into priestly families. Remarkably, many priests preserve family genealogies that can be traced back to the purported time of arrival of Iranian Zoroastrians in India, and beyond to an Iranian homeland.

Genetic data provide a means of examining the biological relationships of different populations and testing claims of common ancestry. Previous studies of Iranian Zoroastrians have suggested they are genetically differentiated from their neighbouring populations. For example, Farjadian et al.^7^ analysed mitochondrial DNA (mtDNA) variation in 14 different ethnic groups from Iran and observed that Zoroastrians and Jews were genetically distinct from other groups. In the same vein, Lashgary et al.^8^ analysed fourteen bi-allelic loci from the non-recombining region of the Y-chromosome (NRY) and observed a notable reduction in haplogroup diversity in Iranian Zoroastrians compared with all other groups. Furthermore, a recent study using genome-wide autosomal DNA found that haplotype patterns in Iranian Zoroastrians matched more than other modern Iranian groups to a high coverage early Neolithic farmer genome from Iran^9^.

Less is known about the genetic landscape and the origins of Zoroastrianism in India, despite Parsis representing more than 80% of present-day Zoroastrians worldwide^10^. A study of four restriction fragment length polymorphisms (RFLP) suggested a closer genetic affinity of Parsis to Southern Europeans than to non-Parsis from Bombay^11^. Furthermore, NRY haplotype analysis^12^ and patterns of variation at the HLA locus^13^ in the Parsis of Pakistan support a predominately Iranian origin of these Parsis.

Prompted by these observations we explored the genetic legacy of Zoroastrianism in more detail by generating novel genome-wide autosomal and Y/mtDNA genotype data for Iranian and Indian Zoroastrian individuals. By comparing to other publicly available genetic data and exploiting linkage disequilibrium information in the autosomal genome, we aimed to identify the demographic processes, including admixture and isolation, that have contributed most to shaping the current genetic landscape of modern Zoroastrian populations. We used the priestly status of Zoroastrian individuals to evaluate claims of patrilineal recent common ancestry. We also assessed the extent to which genetic data supports historical records tracing the origin of Indian Zoroastrians to migrants from Iran, including the timing of migrations and the patrilineal and matrilineal contributions of Iranian Zoroastrians to the Parsi gene pool. Finally, we searched for genomic signatures of positive selection in the Zoroastrian populations that may relate to the prevalence of diseases or other phenotypic traits in the community.

## Materials and methods

### Samples

Buccal swabs were collected from a total of 526 men from India, Iran, the United Arab Emirates and the United Kingdom (see Table S1). Individuals sampled in the United Arab Emirates are mainly first generation Parsis who left Aden following the communist coup in 1970, after which Asians were expelled (Aden was part of the Bombay Presidency until 1947 and the British left Aden in 1967-1968). Individuals sampled from the United Kingdom Zoroastrian population are mainly descendants of 19^th^ century immigrants; the Zoroastrian Association was formed in 1861 at which time there were around 50 Zoroastrians living in the UK^14^. Swabs were stored in a DNA preservative solution containing 0.5% Sodium Doecyl Sulphate and 0.05 M EDTA for transport purposes and DNA was purified by phenol-chloroform extraction/isopropanol precipitation. Informed consent was obtained from all individuals before samples were taken.

### Genome-wide genotyping with the Human Origins array

71 of these samples (29 Iranian Zoroastrians, 17 Iranian Fars, 13 Indian Zoroastrians and 12 Indian Hindu) all of them belonging to the lay (i.e. non-priest) population were genotyped using the Affymetrix Human Origins array, which targets 627,421 Single-Nucleotide-Polymorphisms (SNPs) with well-documented ascertainment, though we note that our techniques here use haplotype information which have been shown to be less affected by ascertainment bias^15,16^. SNPs and individuals were pruned to have genotyping rate greater than 0.95 using PLINK v1.9^17^. Genotypes for the Iranian Zoroastrians and the Iranian Fars were made publicly available by Broushaki et al. ^9^

The above mentioned dataset was then merged with modern populations in the Human Origins dataset of Lazaridis et al.^18^, which includes 17 labelled populations from India and Iran. We also included other high coverage ancient samples: the early Neolithic WC1^9^, Mesolithic hunter-gatherer from Luxembourg (Loschbour), Neolithic individuals from Germany (LBK), Anatolia (Bar8^19^), Georgia (KK1^20^), and Hungary (NE1^21^), a 4,500 year old genome from Ethiopia (Mota^22^) and 45,000 year old genome from western Siberia Ust-Ishim^23^. In total, the merge contained 2,553 individuals and 525,796 overlapping SNPs.

### Principal Component Analysis (PCA)

We performed PCA on all the South Asian and West European populations included in the merge using PLINK 1.9 after LD pruning using –indep-pairwise 50 5 0.5.

### Phasing

We jointly phased the autosomal chromosomes for all individuals in the merge using SHAPEIT^24^ with default parameters and the linkage disequilibrium-based genetic map build 37.

### Chromosome painting and fineSTRUCTURE

We classified our 2545 modern individuals into 230 groups, with the majority of these groups based on population labels^18^. The exceptions to this are the individuals from Iran and India and neighbouring populations of interest for this work (originally labelled as: Onge, Mala, Tiwari, Kharia, Lodhi, Vishwabrahmin, GujaratiD_GIH, GujaratiB_GIH, GujaratiA_GIH, GujaratiC_GIH, Cochin_Jew, India_Hindu, India_Zoroastrian, Iranian, Iran_Fars, Iran_Zoroastrian, Iranian_Bandari, Iranian_GM, Iranian_Shi, Iranian_Lor, Iranian_Jew, Brahui, Balochi, Hazara, Makrani, Sindhi, Pathan, Kalash, Burusho, Punjabi_Lahore_PJL, Druze, BedouinB, BedouinA, Palestinian, Syrian, Lebanese, Jordanian, Yemen, Georgian_Megrels, Abkhasian, Armenian, Lebanese_Christian, Lebanese_Muslim, Assyrian, Yemenite_Jew, Turkish_Jew, Turkish_Kayseri, Turkish_Balikesir, Turkish, Turkish_Istanbul, Turkish_Adana, Turkish_Trabzon, Turkish_Aydin, Iraqi_Jew, Georgian_Jew, AltaiNea, DenisovaPinky, UstIshim, GB20, KK1, LBK, Loschbour, NE1, Bar8, WC1). Individuals from these groups were re-classified into new, label-independent groups using results from the genetic clustering algorithm fineSTRUCTURE that groups individuals into genetically homogeneous clusters based entirely on patterns of shared ancestry identified by CHROMOPAINTER^25^. Briefly, CHROMOPAINTER uses a “chromosome painting” approach that compares patterns of haplotype sharing between each recipient chromosome and a set of donor chromosomes^25^. For the CHROMOPAINTER analysis used for our fineSTRUTURE analysis, which is the first painting protocol described in this paper and referred to throughout as the “fineSTRUCTURE painting”, we painted each of the 696 individuals from the 65 populations listed above using all other 695 individuals as donors. Here we initially estimated the mutation/emission (Mut, “-M”) and switch rate (Ne, “-n”) parameters using 10 steps of the Expectation-Maximisation (E-M) algorithm, for chromosomes 1, 4, 15 and 22, and for every 10 individuals, which gave estimated Mut and Ne of 0.00091 and 320.9197, respectively. These values were then fixed before running CHROMOPAINTER across all chromosomes to produce a “painting profile” giving the proportion of genome wide DNA each individual shares with each other donor individual in this analysis. All chromosomes were then combined to estimate the fineSTRUCTURE normalisation parameter “c”, which was 0.279452. Following Leslie et al.^26^, we then ran fineSTRUCTURE using this “c” value and performing 2,000,000 iterations of Markov-Chain-Monte-Carlo (MCMC), sampling an inferred clustering every 10,000 iterations. Following the recommended approach described by Lawson et al.^25^, we next used fineSTRUCTURE to find the single MCMC sampled clustering with highest posterior probability and performed 100,000 additional hill-climbing steps to find a nearby state with even higher posterior probability. This hill-climbing approach grouped these 695 individuals into 207 clusters, which we then merged into a tree using fineSTRUCTURE’s greedy algorithm that merges pairs of clusters, step at a time, until only two super-clusters remain.

Based on this tree and visual inspection of haplotype sharing patterns among our 207 clusters, we classified these clusters into genetically homogeneous groups, choosing a level of the tree where there were K=50 total clusters. At this level of the tree, we note that the 10,000-year-old Neolithic Iranian WC1 clustered with other modern Iranians, but nonetheless we re-classified WC1 as its own cluster, so that we ended up with 51 final total clusters we use throughout this paper (see Table S2, Figure S1). One of these 51 clusters contained all 13 Indian Zoroastrians or Parsis and represents the “Parsis” group we use throughout this paper. A separate cluster contained 27 of 29 Iranian Zoroastrians plus a single Fars individual that was very genetically similar to self-identified Zoroastrians (Figure S2, Table S2). This particular Fars individual (IREJ-T053) was collected in the city of Yazd, home to one of the oldest Zoroastrian communities in Iran, so it is plausible that this individual might have been mislabelled or recently converted from Zoroastrianism to Islam. Hence we did not remove this Fars person, and instead used all 28 individuals (i.e. the 27 Iranian Zoroastrians plus this Fars individual) to represent the “Iranian Zoroastrian” group we use throughout this paper.

We then painted all 230 modern and 8 ancient samples using all 230 modern groups as donors, following the “leave-one-out” approach, as described by Hellenthal et al.^27^, which is designed to make the final painting profiles comparable. In particular if each donor group {1, …, K} contains {n_1_, …, n_K_} individuals, respectively, the set of donors is fixed to contain n_k_ – 1 individuals from each of the K groups. This is to account for the fact that individuals cannot be painted using themselves as a donor, so that individuals within each of these K donor groups can only ever be painted using n_k_ – 1 individuals from their own group label. We refer to this second painting protocol where K=230 as the “all donors painting” throughout. Note that a primary difference between this painting and the “fineSTRUCTURE painting” described above is that we now use group labels, based in part on clustering results, which are required for our leave-one-out approach. When using haplotype information for this painting, we initially estimated the mutation/emission and switch rate parameters as described above, giving estimated Mut and Ne of 0.000704 and 223.5674, respectively. Alternatively, where noted below we used an “unlinked” approach (-u switch) that analysed each SNP independently (i.e. ignored haplotype information) using the default CHROMOPAINTER emission rate.

We also performed a slightly different version of this painting where Iranian and Indian populations were excluded as donors, using the leave-one-out approach described above for all 216 other groups, a third painting protocol with K=216 that we refer to throughout as the “non Indian/Iranian donors painting”. We did this to infer how Iranian and Indian groups relate ancestrally to groups from outside their own countries, which for example can help determine whether admixture from outside groups (rather than independent drift effects due to genetic isolation) is driving genetic differences among these sampled groups within Iran and India^26,28^. Mut and Ne parameters (0.00069 and 225.32, respectively) were re-estimated for this new scenario as described above. When painting Iran, we excluded all Iranian and Indian individuals as donors. In contrast, we painted the Indian groups using all non-Indian groups as donors – i.e. we included Iranian groups as donors. This is because in this paper we infer Iranians as important contributors to the DNA of Parsis, making them important to include when evaluating genetic differences among Indian groups that are due to admixture.

### TVD, F_XY_ and F_ST_ between Iranian and Indian groups

We quantified differences in the painting profiles between all Iranian and Indian groups by applying the metric total variation distance (TVD) as described in Leslie et al.^26^ using the formula:

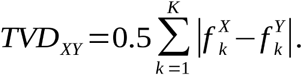

where 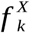 and 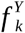 are the average genome-wide proportion of DNA that individuals from the recipient groups X and Y, respectively, match to donor group *k* ∈ [1, …, *K*] as inferred by CHROMOPAINTER. For this paper TVD was calculated using the “all donors painting” results from runs of CHROMOPAINTER that a) used haplotype information and b) used an “unlinked” approach that ignores haplotype information and instead analyses all SNPs independently.

Independent drift effects in groups X and Y since their split can generate genetic differences between them without requiring any outside introgression since this split. To elucidate whether inferred genetic differences, e.g. as measured by are more attributable to ancestry from outside sources, we followed the approach in van Dorp et al.^28^ designed to mitigate these drift effects. In particular we calculated:

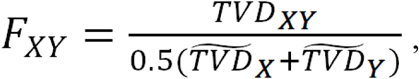

where 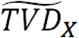 equals

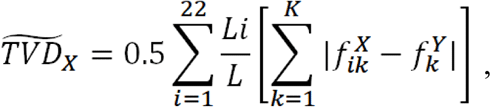

where L_i_ is the number of SNPs in chromosome i ∈ [1,…,22], L the total number of SNPs across all the 22 chromosomes, and 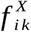 is the average proportion of DNA that individuals from X match to donor group k when painting only chromosome i. This approach scales genetic differences between the two groups by differences across chromosomes within each group, exploiting how each chromosome should be subjected to independent drift^29^. For this analysis we used the “Non-Indian/Iranian donors painting” that excluded Indian and Iranian populations as donors in the dataset, which similarly attempts to attenuate drift effects within each Iran and Indian group by matching their DNA to only groups outside of their countries (thus disallowing “self-copying” in Iran/Indian groups)^28^. For comparison purposes, F_XY_ was also calculated for the Iranian and Indian groups using the “all donors painting” (Figures S4-S5).

The weighted F_ST_ for these groups was also calculated based on independent SNPs using PLINK 1.9, which implements the method introduced by Weir and Cockerham^30^.

### Exploring relative amounts of genetic diversity within groups

For a comparison of techniques, we used the following three distinct approaches to quantify the relative amounts of genetic diversity within groups:

#### (1) CHROMOPAINTER analyses to infer relative amounts of genetic diversity within groups

We performed a fourth analyses using CHROMOPAINTER that is analogous to that in van Dorp et al.^28^, to assess the relative genetic diversity within our 8 fineSTRUCTURE inferred clusters with sample size greater than or equal to 13, which is the number of Parsis individuals: Indian_A, Indian_B, Indian_C, Parsis, Iranian_A, Iranian_Zoroastrian, Kharia, Mala_Vishwabrahmin. For each of these 8 clusters, we randomly subsampled 13 individuals and painted each individual using only the other 12 individuals from their respective cluster as donors, using 50 steps of CHROMOPAINTER E-M algorithm inferring the switch and emission rates (i.e. “-i 50 -in -iM”). We refer to this fourth painting protocol throughout as the “within-group-diversity painting”. For each individual, we calculated average segment size by dividing the total proportion of genome-wide DNA copied from all donors by the total expected number of haplotype segments copied from all donors. This average segment size can be thought of as capturing the relative amount of genome-wide haplotype diversity in each group, with a relatively larger average segment size reflecting relatively less genome-wide diversity.

#### (2) PLINK IBD analysis to infer relative amounts of genetic diversity within groups

We also inferred within group genetic diversity across all pairwise combinations of individuals within each of the above genetic clusters (Indian_A, Indian_B, Indian_C, Parsis, Iranian_A, Iranian_Zoroastrian, Kharia, Mala_Vishwabrahmin) using the IBD coefficient PI_HAT implemented in PLINK v1.9, on a dataset where SNPs were first pruned to remove those in high linkage disequilibrium (r^2^>0.2) in a sliding window of 250 SNPs. For consistency, the same 13 individuals were used to calculate the genetic diversity within each group based on PI_HAT as in calculating haplotype segment size.

#### (3) fastIBD analysis to infer relative amounts of genetic diversity within groups

In order to explore within group genetic diversity using a third approach, which allows SNPs to be in LD as in our CHROMOPAINTER-based estimates of segment size, we applied fastIBD using the software BEAGLE v3.3.2^31^. For each cluster, we used the same subset of 13 individuals randomly sampled above and used fastIBD to infer the pairwise IBD fraction between each pairing of these individuals. For each chromosome of each cluster, fastIBD was run for 10 independent runs and an IBD threshold of 10^−10^ for every pairwise comparison of individuals as recommended by Browning and Browning^31^ though we note results were similar using an IBD threshold of 0.0001.

### Inferring admixture events using a mixture modelling approach, GLOBETROTTER, f3-statistics and TreeMix

As noted previously^26,27^, the inferred CHROMOPAINTER painting profiles are often not the best summary of shared ancestry patterns, as for example donor groups with larger sample sizes may be disproportionately represented in these paintings. In order to account for this we performed additional analyses to “clean” the raw CHROMOPAINTER output. In particular, we applied the Bayesian mixture modelling approach described in Broushaki et al.^9^ to infer proportions of ancestry for all recipient groups (which we term “targets”) in relation to other included groups that represent potential “surrogates” to sources of ancestry. Here we performed two analyses: (a) including all 229 modern groups excluding the target as potential surrogates and (b) using all 229 modern and 8 ancient groups as potential surrogates (i.e. 237 surrogate groups in total). The aim of this mixture modelling approach is to identify which subset of these 229-237 potential surrogates best reflect the sources of ancestry in the target group. We then use this subset of surrogates in our admixture analysis described below. However, we note that any inferred proportions from this mixture model analysis cannot necessarily be interpreted as reflecting proportions of admixture from distinct source groups. Instead this mixture modelling step is primarily used to summarize the clearest patterns of shared ancestry between the target and surrogate groups, and to restrict the set of surrogates used in our subsequent admixture analysis to help increase power and precision.

We applied GLOBETROTTER^27^, a haplotype-based approach to identify, describe and date recent admixture events, to test for evidence of admixture separately in each of 24 “target” groups from Iran, India, Pakistan and Armenia. Roughly speaking, GLOBETROTTER infers admixture in a target group using two (interlocking) steps. The first infers the genetic make-up of the putative admixing source groups, and the second infers the date of admixture. For the first step we used the “all donors painting” from CHROMOPAINTER for each target group, as this GLOBETROTTER inference step requires each surrogate and target group to be painted using the same (or a very similar) set of donors^27^. While for the second step, we used CHROMOPAINTER to generate 10 painting samples per haploid genome for each Iranian, Indian, Pakistani and Armenian individual, under a different painting where each of these individuals is painted excluding any individuals from their assigned group as donors. We refer to this a fifth painting protocol as “GLOBETROTTER painting”. We do this fifth “GLOBETROTTER painting” to follow the suggested protocol in Hellenthal et al.^27^, as including individuals from your own group as donors when painting often substantially masks signals of admixture, particularly when generating the linkage disequilibrium (LD) decay curves critical to dating admixture. This is because individuals (unsurprisingly, but unhelpfully) match large segments of their genome to other individuals from their own group. While we could also use this “GLOBETROTTER painting” for the first step that infers the genetic make-up of the admixing source groups, for each target group we would then have had to re-paint every surrogate group similarly excluding that target group’s individuals as donors. For computational simplicity we instead used the same “all donors” painting for each target group, which previous work suggests makes little difference in practice for these sample sizes and which we explore further below^27^. For each target population we included only the surrogate groups that contributed to our mixture modelling approach described above, separately under the two mixture modelling scenarios using as surrogates (a) modern groups only and (b) modern and ancient groups. We inferred admixture dates using the default LD decay curve range of 1-50cM and bin size of 0.1cM when considering the distance between genome segments. An exception to this is cases where the inferred admixture date was >60 generations ago using this default curve range and bin size, in which case we re-estimated dates using a curve range of 1-10cM and a bin size of 0.05cM, as this has been shown previously to more reliably estimate older dates of admixture^27^, In each analysis we used 5 iterations of GLOBETROTTER’s alternating source composition and admixture date inference (num.mixing.iterations: 5) and 100 bootstrap re-samples to infer confidence intervals around the point estimates of the date of admixture. Furthermore, in each case analyses were run twice, once using the option null.ind:0 and once with null.ind:1 to assess the effect of standardizing against a pseudo (null) individual, an approach designed to account for spurious signals of linkage disequilibrium that are not attributable to admixture^27^. Only results for null.ind =1 are shown, as results for null.ind=0 were very consistent. For comparison, we also performed an additional GLOBETROTTER analysis using the surrogates inferred under (a) and (b) when using CHROMOPAINTER results from the “non Indian/Iranian donors painting”, this time using the same CHROMOPAINTER painting for both the first and second steps of GLOBETROTTER described above.

As a very different means of inferring admixture, we also used ADMIXTOOLS^29^ to calculate f3 statistics, f3(X; A,B), a commonly-used test to detect admixture in a target population X presumed to have received DNA from two ancestral source populations represented by surrogate groups A and B. We inferred admixture separately in the Indian and Iranian Zoroastrians, using all pairwise combinations of the other populations in the dataset, plus the ancient samples, as possible admixture sources A and B.

Additionally, we used TreeMix^32^ to infer a bifurcating tree that merges four groups: our Indian and Iranian Zoroastrian groups, and the groups with largest sample size from each of Iran and India. We also included the Yoruba as the outgroup (root) population, allowed different numbers of migration events (0-3) among populations in the tree, and accounted for linkage disequilibrium between SNPs grouping them in windows of 500 SNPs (-k 500).

### Positive Selection tests

We used the XP-EHH (Cross Population Extended Haplotype Homozygosity) statistic^33^ to detect signatures of recent positive selection by comparing populations with similar demographic histories. Thus, we inferred putative regions of positive selection in Zoroastrians of Iran and India, using as reference populations the clusters Iranian_A and Indian_A (for the latter only the individuals labelled as India_Hindu and Gujarati were used due to usage restriction of the other samples for selection tests^18^), respectively. Normalized XP-EHH scores were calculated using SELSCAN v.1.1.0^34^. The direction of selection was determined by the sign of the XP-EHH scores, with positive values indicating selection in the Zoroastrian populations and negative values indicating selection in the non-Zoroastrian populations. SNP annotations were obtained using ANNOVAR^35^. Here we apply XP-EHH to populations we infer to be admixed (see Results). While XP-EHH has been applied to admixed populations before^36^, we note this presumably may lead to spurious findings, as proportions of DNA inherited from an introgressing group (which may have more or less linkage disequilibrium than the ancestral group) will vary across genetic regions.

To assess the significance threshold of the analysis, we performed 100 permutation tests to establish the empirical distributions of XP-EHH values across the genome for both the Indian and Iranian populations. For each permutation, we randomly partitioned our Zoroastrians and non-Zoroastrians into [two different groups, and then calculated XP-EHH comparing these two groups. The threshold values at significance level of 0.01% (quantiles 0.0001 and 0.9999) from the empirical distribution combining all 100 permutations were used to determine the significance of the XP-EHH test. These values were of -4.46 and 4.46 for the Iran, and -4.37 and 4.37 for India.

### Non-recombining region of the Y-chromosome (NRY) and mitochondrial DNA (mtDNA) analysis using data from the Human Origins array

NRY haplogroups were assigned to Indian (India_Hindu, Mala, Tiwari, Vishwabrahmin and India_Zoroastrian), Pakistani (Balochi, Brahui, Burusho, Hazara, Kalash, Kharia, Makrani, Pathan and Sindhi) and Iranian (Iranian and Iran_Fars jointly analysed, and Iran_Zoroastrian) populations from this dataset using a maximum likelihood approach against the Y-chromosome consortium NRY phylogenetic tree^37^ with Yfitter^38^. Individuals for which NRY haplogroup could not be assigned to were removed from further analysis. Individuals were assigned to known mitochondrial haplogroups based on observed mtDNA SNP variation with HaploGrep^39^. F_ST_ genetic distances^40^ were estimated among all the groups based on NRY or mtDNA haplogroup frequencies using Arlequin version 3.1^41^.

### Additional Y-chromosome typing and mitochondrial DNA sequencing

In order to further explore sex biased admixture and to evaluate claims of patrilineal inheritance among the Parsi priests, all the 526 samples collected for this study were typed for Y-chromosome (490 successful samples) and mitochondrial DNA (518 successful sequencing) (see Table S1). Y-chromosomes were typed for six STRs (DYS19, DYS388, DYS390, DYS391, DYS392, DYS393) and at 11 biallelic loci (92R7, M9, M13, M17, M20, SRY1465, SRY4064, SRY10831, sY81, Tat, YAP) as described by Thomas, Bradman, and Flinn^42^, and for the biallelic marker 12f2 as described by Rosser et al.^43^ Microsatellite repeat numbers were assigned according to the nomenclature of Kayser et al.^44^ For a subset of the samples (Parsi priests), four additional Y-chromosome microsatellites (DYS389I, DYS389II, DYS425 and DYS426) were typed as described by Thomas, Bradman, and Flinn^42^. Y-chromosome haplogroups (Yhg) were defined by the 12 biallelic markers according to a nomenclature modified from Rosser et al.^43^ and Weale et al.^45^ The correspondence between this nomenclature and that proposed by the YChromosome Consortium^46^ is as follows: Yhg-1 = 5 P*(xR1a), Yhg-2 = 5 BR*(xDE,JR), Yhg-3 = 5 R1a1, Yhg-4 = 5 DE*(xE), Yhg-7 = 5 A3b2, Yhg-8 = 5 E3a, Yhg-9 = 5 J, Yhg-16 = 5 N3, Yhg-20 = 5 O2b, Yhg-21 = 5 E*(xE3a), Yhg-26 = 5 K*(xL,N3,O2b,P), Yhg-28 = 5 L, Yhg-29 = 5 R1a*, Yhg-37 = 5 Y*(xBR,A3b2).

The mitochondrial DNA hyper variable segment 1 (HVS-1) was sequenced as described by Thomas et al.^47^ Sequences were obtained for all samples between positions 16,027 and 16,400 according to the numbering scheme of Anderson et al.^48^ MtDNA haplotypes were assigned to haplogroups (iMhg) firstly by identifying key combinations of HVS-1 alleles according to Macaulay et al.^49^, Richards et al.^50^ and Maca-Meyer et al.^51^ as follows: 16129A, 16223T, 16391A = iMhg-I, 16069T, 16126C = iMhg-J, 16224C, 16311C = iMhg-K, 16126C, 16294T = iMhg-T, 16223C, 16249C = iMhg-U1, 16223C, 16051G = iMhg-U2, 16223C, 16343G = iMhg-U3, 16223C, 16356C = iMhg-U4, 16223C, 16270T = iMhg-U5, 16223C, 16318T = iMhg-U7, 16223T, 16292T = iMhg-W, 16189C, 16223C, 16278T = iMhg-X. For the remaining haplotypes, those with a T at position 16223 were assigned to iMhg-MNL and those with a C at position 16223 were assigned to iMhg-HVR.

Unbiased genetic diversity, *h,* and its standard error were calculated using the formula given by Nei^52^ and significant differences in calculated values were found using a standard two-tailed *z* test. Populations were compared using F_ST_ based on haplotype or haplogroup frequencies, estimated from analysis of molecular variance (AMOVA) Ø_ST_ values^40,53^, and using the Exact Test for Population Differentiation^54^. Assessment of the significance of pairwise F_ST_ values was based on 10,000 permutations of the data and 10,000 Markov steps were used in the Exact Test. Patterns of genetic differentiation were visualized using principal coordinates (PCO) analysis performed on a similarity matrix calculated as one minus F_ST_, based on Yhg and iMhg frequencies. Values along the main diagonal of the similarity matrix, representing the similarity of each population sample to itself, were calculated from the estimated genetic distance between two copies of the same population sample (for Ø_ST_ -based F_ST_, the resulting self-similarity values simplify to *n*/(*n* - 1), where *n* is the sample size).

Y-chromosome and mtDNA admixture proportions were estimated using the likelihood-based method LEA^55^, based on Yhg and inferred iMhg frequencies. We ran 5,000,000 Monte Carlo iterations of the coalescent simulation and discarded the first 10,000 iterations as burn-in. For comparison admixture proportions mY, mC and mR were also estimated, using the methods of Bertorelle and Excoffier^56^, Chakraborty et al.^57^ and Roberts and Hiorns^58^ respectively. 10,000 bootstrap re-samplings were carried out to estimate standard errors and admixture proportions were compared using a standard two-tailed *z* test.

The coalescence time of clusters of Y-chromosomes belonging to the same UEP defined haplogroup was estimated by finding the Average Square Difference (ASD) between the inferred ancestral haplotype (in this case the modal haplotype) and all observed chromosomes^59,60^. The 95% confidence interval for this estimate was calculated as described in Thomas et al.^61^ using 50,000 iterations. The microsatellite mutation rate was set to 15/7856, based on data from three published studies^62,63,64^. This analysis was restricted to haplogroups containing a high frequency modal haplotypes (>50%) where the ancestral state could be inferred with confidence.

## Results

### Zoroastrians are genetically differentiated from non-Zoroastrians, with different historical ancestry in Parsis relative to non-Zoroastrian Indians

Most of the Iranian Zoroastrians (see Methods and Table S1 for a description of the samples used in this work) are positioned within the autosomal genetic variation of other sampled Iranian samples in a PCA of West Eurasian individuals (Figure S3). Interestingly, two of the 29 Iranian Zoroastrians (YZ020 and YZ024) look genetically different from the others, and were inferred by fineSTRUCTURE to cluster with other non-Zoroastrian Iranians (Figures S1-S2), which is consistent with Zoroastrians not being as closed a community as is sometimes thought and reported^6^. We will come back to this issue later, but in order to study the common ancestry of the genetically homogeneous majority of our sampled Iranian Zoroastrians, these two individuals were excluded from further analysis. The Parsis (i.e. Indian Zoroastrians) form a more wide-ranging cluster along PC1, falling inside Iranian, Pakistani and Indian groups (Figure S3).

We clustered some of our sampled individuals, including all Indians and Iranians, into 51 genetically homogeneous groups that exhibited good correlation between genetic similarity and population label (Figure 1a, Figure S1, Table S2; see Methods for explanation of clustering approach). One of these 51 clusters contained all 13 Parsis, forming the “Parsis” group we use throughout the remainder of this study. A separate cluster contained 27 of 29 Iranian Zoroastrians plus a single Farsi individual that was very genetically similar to self-identified Zoroastrians (Figure S2, Table S2), and these 28 individuals form the “Iranian Zoroastrian” group we use throughout the remainder of this study. The remaining genetically homogeneous clusters (Figure 1a, Figure S1, Table S2) containing Indian and Iranian individuals that we refer to below consist of: (1) 54 Iranians primarily from Lori, Shiraz and Yazd and Iranian Mazanderanis (referred to as “Iranian_A”); (2) 7 Bandari Iranians (“Iranian_B”); (3) 2 genetically distinct Bandari Iranians (“Iranian_C”); (4) 7 Iranian Jews (“Iranian_Jews”); (5) 16 primarily Gujarati Indians (“Indian_A”); (6) 24 Tiwari and Gujarati Indians (“Indian_B”); (7) 25 Lodhi and Hindu Indians (“Indian_C”); (8) 26 Mala and Vishwabrahmin Indians (“Mala_Vishwabrahmin”); (9) 13 Kharia Indians (“Kharia”); (10) 11 Onge (“Onge”); (11) 2 Cochin Jews from India (“Cochin Jews_A”); and (12) 3 other Cochin Jews from India (“Cochin Jews_B”) (Figure 1, Table S2).

**Figure 1.**
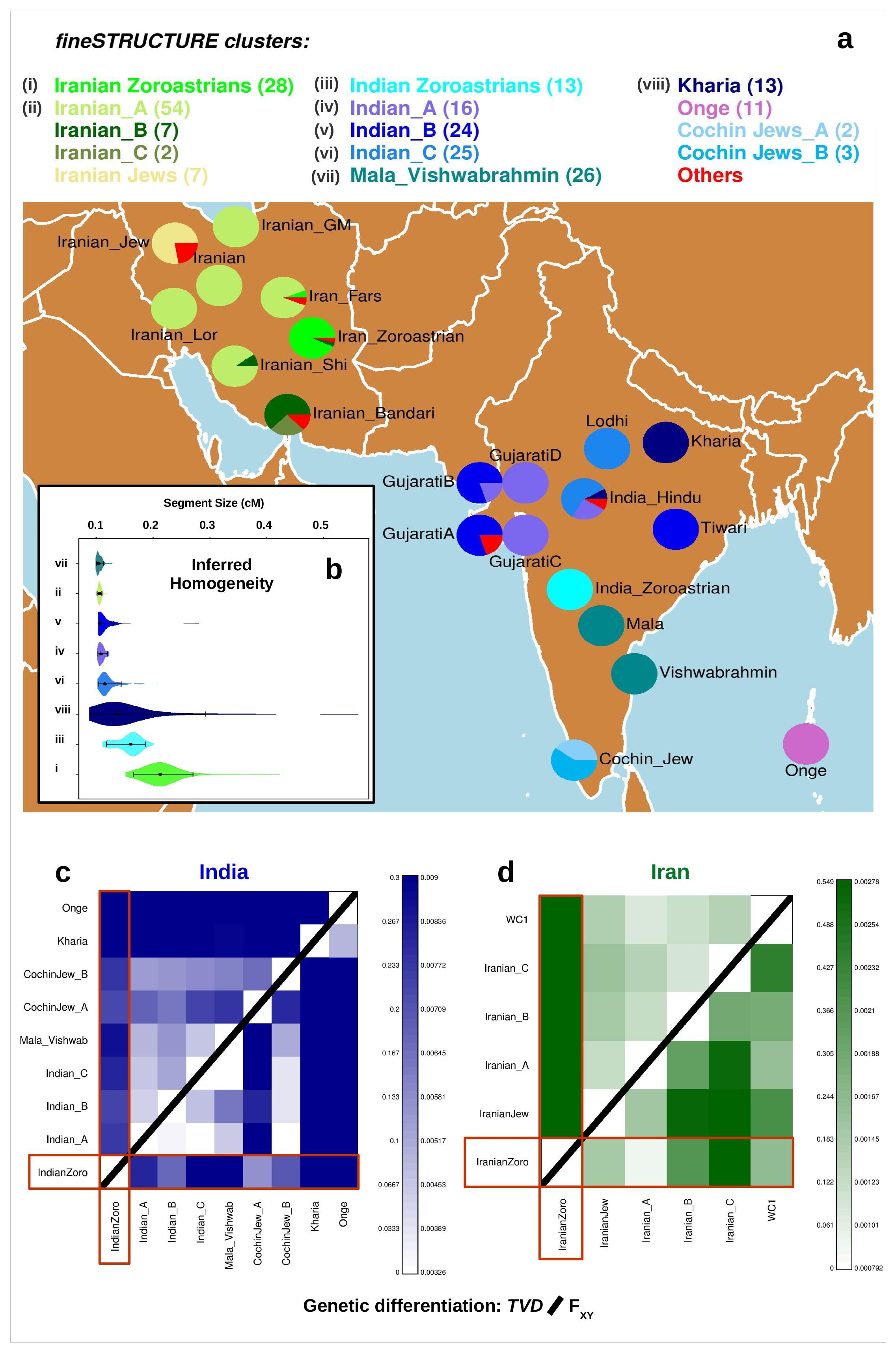
Clustering, homogeneity and genetic differentiation of the Iranian and Indian populations. (a) Each color inside the pies represents the proportion of individuals from each population label that is assigned to each fineSTRUCTURE cluster (“Others” include all groups outside Iran and India), with the total number of individuals included in each cluster shown inside brackets in the legend. (b) Distribution of CHROMOPAINTER’s inferred lengths of haplotype segments (in cM) copied intact from a single donor, when allowing 13 randomly-sampled individuals from each group (roman numerals in part (a) legend) to copy from the other 12 individuals with the same label. (Black dot = median values, bars = 95% empirical quantiles across individuals.) (c)-(d) Comparison of pairwise TVD based on the “all donors painting” (upper triangle) and FXY based on the “non-Indian/Iranian donors painting” mitigating recent drift effects (lower triangle) for (c) Indian and (d) Iranian groups.

Among our sampled individuals from Armenia, India, Iran and Pakistan, we measured genetic distance between pairs of groups using two different techniques: (1) the commonly-used, allele frequency-based measure F_ST_^30^, and the haplotype-based measure (2) TVD^26^ (see Methods; Table S3, Figures S4-S5). While genetic distance among groups is not large overall (e.g. typically F_ST_ < 0.04), similar to Jewish groups from these regions, the Onge from the isolated Andaman Islands, and the Indian Kharia, an indigenous tribal ethnic group that has been isolated from other groups^65^, Zoroastrians were strongly genetically differentiated from non-Zoroastrians under each of these three measures, agreeing with previous work^7,8,66^. For example, the genetic distance between Iranian Zoroastrians and non-Jewish, non-Zoroastrians from Iran ranged from 0.015-0.029 for F_ST_ and 0.544-0.551 for TVD, with each distance measure larger than the maximum such measure between any two non-Zoroastrian, non-Jewish Iranian groups (0.011 and 0.164, respectively) (Table S3). Similarly, excluding the Onge and Kharia, the genetic distances between Parsis and non-Zoroastrians from India ranged from 0.014-0.028 for F_ST_ and 0.221-0.278 and TVD, with each measure larger than the maximum distance between any two other non-Zoroastrian Indian groups (0.002-0.008 and 0.058-0.122, respectively). Therefore, in both Iran and India, these results indicate a high degree of genetic distance between the Zoroastrians in these countries relative to most other sampled individuals from their respective countries.

Our haplotype-based techniques are designed to identify which sampled individuals share ancestors with each other most recently. Typically, individuals share more recent ancestors with individuals of the same population label than with individuals from other populations, as is the case here with both Zoroastrian groups, reflecting (sometimes recent) genetic isolation between individuals with different population labels. However, we also measured genetic distance between pairs of groups using a different haplotype-based genetic distance measure, F_XY_, and an analysis (“Non Indian/Iranian donors painting”; see Methods) that was specifically designed to mitigate signals of genetic differentiation attributable to recent genetic isolation^28^. Briefly, we do this by comparing the DNA of individuals from a particular group only to other individuals that were sampled from other geographic areas, for example comparing the DNA of Iranian Zoroastrians to only that of non-Iranians and non-Indians. Relative to many of the ancestors shared among people from the same country, this inference often reflects sharing of ancestors that lived farther back in time. In practice this painting and our F_XY_ score, which uses independent drift effects across chromosomes to further substract our genetic differentiation due to recent isolation, should indicate a relatively small amount of genetic distance between two groups that have a similar recent ancestral history, e.g. have similar sources of admixture from outside sources or descend from a common recent source population. This should be true even if the two groups have largely stopped intermixing with one another for a period of time, such that they have e.g. relatively high F_XY_ and TVD^28^. Under this F_XY_ measure, Iranian Zoroastrians showed a much-reduced genetic distance to other Iranian groups (Figure 1d), e.g. with Zoroastrians and the Iranian_A cluster having the lowest F_XY_ value out of all comparisons of Iranian groups (Table S3). In contrast to results using our F_ST_ and TVD measures, genetic dissimilarities measured by F_XY_ among the other Iranian groups (Iranian_Jews, Iranian_A, Iranian_B, Iranian_C) are higher, which we explore further below. However, the F_XY_ scores are not noticeably lower between the Parsis and non-Zoroastrian groups from India, with in general the Parsis showing a similar relatively high amount of genetic differentiation as the Kharia, Onge and Indian Cochin Jewish groups to all other Indian groups (Table S3), mimicking our results when comparing these groups using F_ST_ and TVD (Figure 1c).

Therefore, these analyses suggest that a large degree of observed genetic differentiation between Zoroastrians and non-Zoroastrians from Iran is primarily attributable to genetic isolation between Zoroastrians and non-Zoroastrians in the country. In contrast, a large degree of observed genetic differentiation between Parsis and non-Zoroastrians from India is attributable to the Parsis having different ancestry than other Indian groups (Figure 1c,d).

### Genetic homogeneity is higher in Zoroastrian groups, consistent with increased endogamy relative to non-Zoroastrians in Iran and India

We performed additional analyses to measure the amount of genetic homogeneity separately within each cluster. Compared to non-Zoroastrian groups, we found that each of Iranian Zoroastrians and Parsis shared relatively longer haplotype segments with members of their own group (Figure 1b, Figure S6, Table 1), reflecting a higher degree of genetic similarity within each Zoroastrian group relative to non-Zoroastrian groups. This is consistent with both Iranian and Indian Zoroastrians being genetically isolated from non-Zoroastrian groups^28^. This is true under two distinct homogeneity estimators that use haplotype information. The first approach FastIBD^31^ compares the DNA of pairwise combinations of a group’s individuals, and here gave median shared haplotype lengths of 0.148 cM and 0.113 cM across pairwise combinations of Iranian Zoroastrians and Parsis, respectively, relative to 0.075 for the third largest value in the Kharia. The second approach CHROMOPAINTER^25^ (under our “within-group-diversity painting”; see Methods) compares the DNA of all of a group’s individuals jointly, and here gave median shared haplotype lengths across individuals of 0.212 cM and 0.161 cM for Iranian Zoroastrians and Parsis, respectively, relative to 0.134 for the third largest value in the Kharia. Conflicting slightly with this, we note that the PI_HAT value from PLINK v1.9^17^, which is based on an alternative technique that ignores haplotype information when measuring homogeneity, infer the Kharia to have more homogeneity than Parsis, giving median values of 0.323 and 0.312 across pairwise combinations of Kharia and Parsis, respectively. This perhaps results from a decreased resolution when not exploiting linkage disequilibrium information, at least when using ascertained SNPs^31^.

**Table 1.** Measuring within group homogeneity: Segment size (CHROMOPAINTER), FIBD and PI_HAT. CHROMOPAINTER’s inferred median haplotype segment sizes (in cM) copied intact from a single donor, when allowing 13 randomly-sampled individuals from each cluster to copy from the other 12 individuals assigned to the same cluster, using 50 steps of Expectation-Maximisation (E-M). IBD values inferred by fastIBD (FIBD) implemented in BEAGLE v3.3.2 using the same 13 randomly-sampled individuals. PI_HAT values inferred by PLINK v1.9 across the same 13 randomly-sampled individuals after sub-sampling SNPs to remove those in high linkage disequilibrium are also reported. Median and empirical quantile values across the 13 individuals are given for each metric for each cluster.

| Cluster | Segment (95% CI) | FIBD (95% CI) | PI_HAT (95% CI) |
| --- | --- | --- | --- |
| **Indian_A** | 0.108<br>(0.103-0.117) | 0.063<br>(0.051-0.078) | 0.302<br>(0.297-0.309) |
| **Indian_B** | 0.106<br>(0.104-0.128) | 0.063<br>(0.052-0.075) | 0.301<br>(0.296-0.308) |
| **Indian_C** | 0.115<br>(0.106-0.14) | 0.06<br>(0.05-0.096) | 0.304<br>(0.299-0.312) |
| **Indian Zoroastrians** | 0.161<br>(0.12-0.184) | 0.113<br>(0.074-0.145) | 0.312<br>(0.299-0.324) |
| **Iranian_A** | 0.106<br>(0.103-0.11) | 0.06<br>(0.047-0.08) | 0.301<br>(0.294-0.306) |
| **Iranian Zoroastrians** | 0.212<br>(0.171-0.271) | 0.148<br>(0.098-0.301) | 0.326<br>(0.31-0.372) |
| **Kharia** | 0.134<br>(0.091-0.223) | 0.075<br>(0.052-0.318) | 0.323<br>(0.311-0.412) |
| **Mala_Vishwabrahmin** | 0.104<br>(0.101-0.112) | 0.061<br>(0.049-0.089) | 0.308<br>(0.301-0.319) |

Consistent with our autosomal DNA results, Y/mtDNA results for these same individuals gave gene diversity values that were significantly lower for Iranian and Indian lay Zoroastrians relative to non-Zoroastrians, for both Y-haplotype frequencies (Tables S4 and S6) and mtDNA haplotype frequencies (Tables S5 and S7).

### Evidence for admixture in Zoroastrian groups with different sources and times using nuclear data

We calculated f3 statistics using autosomal DNA from the Iranian Zoroastrians and Parsis as targets and all pairwise combinations of the other modern and ancient groups as sources, reporting all pairwise combinations that gave a negative f3 value with a Z score >|2| for the Parsis in Table S8. In all cases one source of admixture is best represented by a modern-day Indian population. The second source is generally represented by an ancient Neolithic sample from Europe or Anatolia, or a modern group close to Iran such as Armenia, Lebanon, or Iraqi_Jews, suggesting an Iranian-like source. In the case of the Iranian Zoroastrians, no admixture events were inferred with any group present in the dataset, consistent with previous reports of f3 statistics sometimes having decreased power to detect admixture in isolated groups with e.g. bottleneck or founder effects^29^.

Additionally, we identified admixture events in both Parsis and Iranian Zoroastrians by first using a mixture modelling approach^9^ to identify the best ancestry surrogates for each target group, and then running the haplotype-based software GLOBETROTTER to date any putative admixture events using only these surrogates (see Methods). In contrast to f3 statistics, GLOBETROTTER infers the decay of linkage disequilibrium among segments inherited from admixing sources, which increases the power to identify admixture and can also be used to date events^27,67^. For each case we used either (a) only modern groups or (b) both ancient and modern groups as possible surrogates. Each of (a) and (b) gave largely corroborating results, e.g. with confidence intervals for dates overlapping when admixture is inferred for the same target group (Figure 2, Figure S7, Tables S9-S11). However, test (b) was sometimes more sensitive as we note below.

**Figure 2.**
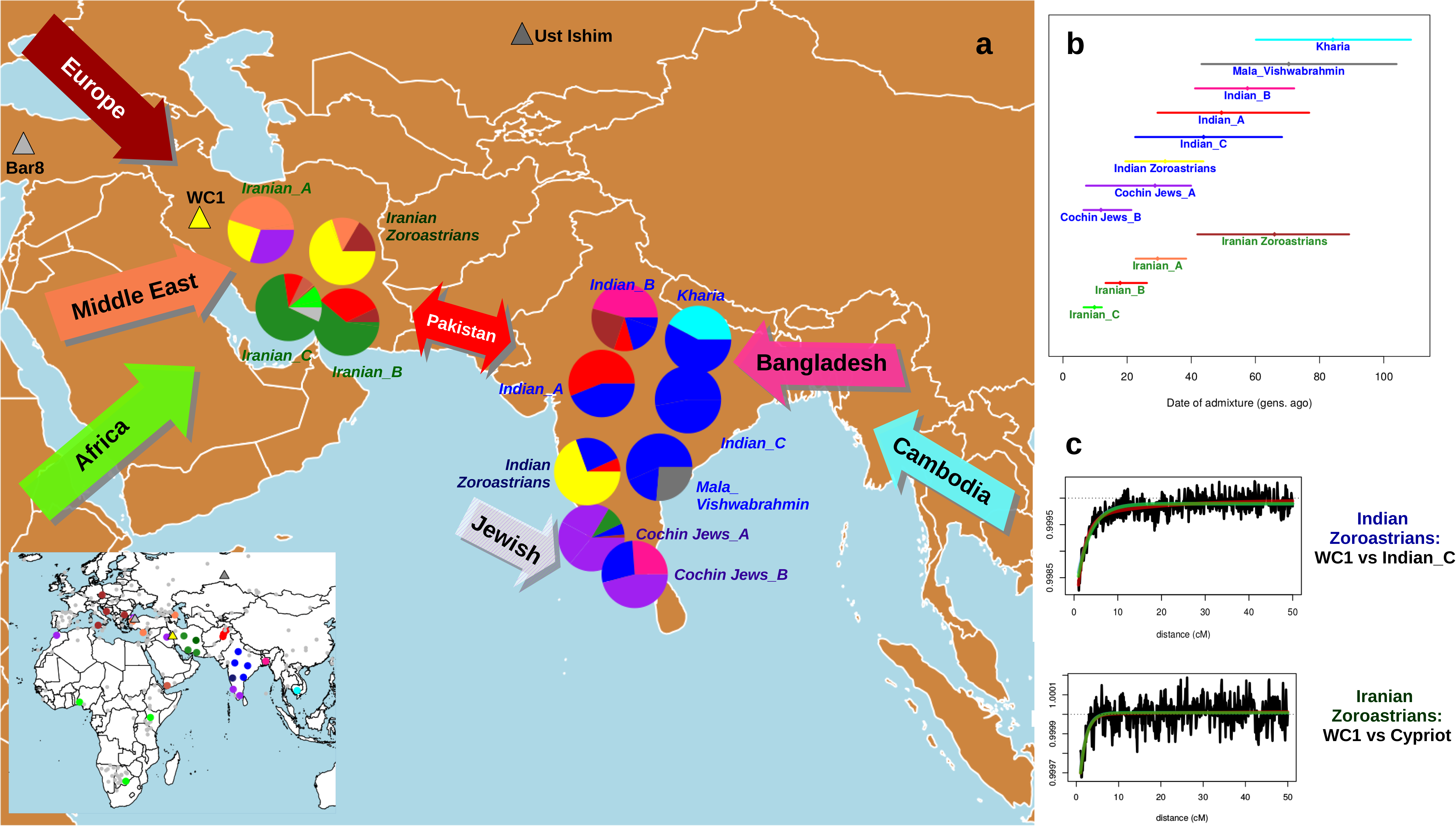
Recent admixture in India and Iran. (a) Inferred recent admixture in India and Iran, using admixture surrogates from Europe (brown), Middle East (orange; Yemen in dark orange), Africa (light green), Pakistan (red), Bangladesh (pink), Cambodia (cyan), Iran (dark green) and India (blue) and of Jewish heritage (purple), plus the ancient samples WC1 (yellow), Ust’Ishim (dark grey) and Bar8 (grey). Inferred proportions of haplotype sharing with each surrogate group are represented in the pie graphs, with all contributing groups highlighted in non-grey in the map in the left bottom corner. (b) Dates of admixture (dots) and 95% confidence intervals (bars) inferred by GLOBETROTTER, colored according to the surrogate that best reflects the minor contributing admixture source. (c) GLOBETROTTER coancestry curves, illustrating the weighted probability (black lines) that DNA segments separated by distance x (in cM) match to the two admixture surrogates given in the title, are given for the Parsis (WC1 vs Indian_C) and Iranian Zoroastrians (WC1 vs Cypriot), along with the best fitting exponential distributions (green lines) using the inferred date from (b) for each.

In (a) and (b) we detected admixture in the Parsis dated to 27 (range: 17-38) and 32 (19-44) generations ago, respectively, in each case between one predominantly Indian-like source and one predominantly Iranian-like source. This large contribution from an Iranian-like source (∼64-76%) is not seen in any of our other 7 Indian clusters, though we detect admixture in each of these 7 groups from wide-ranging sources related to modern day individuals from Bangladesh, Cambodia, Europe, Pakistan, or of Jewish heritage (Figure 2; Figure S7; Tables S9-S11). For Iranian Zoroastrians, we only detect admixture under analysis (b), occurring 66 (42-89) generations ago between a source best genetically explained as a mixture of modern-day Croatian and Cypriot samples, and a second source matching to the Neolithic Iranian farmer WC1. We infer admixture in all three other non-Jewish Iranian groups, though consistently more recent (<38 generations ago) with contributions from sources related to modern-day groups from Pakistan, Sub-Saharan African or Turkey (Figure 2, Figure S7; Tables S9-S11).

We also ran TreeMix on our two Zoroastrian groups, one other Indian group (Indian_C), and one other Iranian group (Iranian_A) in order to infer a bifurcating tree relating these groups, using Yoruba as an outgroup and allowing for 0-3 migration events (Figure S8). While all TREEMIX analyses inferred the highest drift value in the Iranian Zoroastrians, in agreement with our analyses described above, the migration results were less clear despite low residuals (Figure S9). For example, when including admixture TREEMIX inferred migration from ancestors of Iran into Yoruba, though this has never been previously suggested, and from Parsis into other Indian groups rather than the other way around. This likely reflects the challenge in accurately identifying and describing admixture events in some cases when not directly measuring the decay of linkage disequilibrium that is expected in genuine admixture signals^27,67^.

### Evidence for sex-biased admixture in Parsis using Y-chromosome and mtDNA data

Analysis of mtDNA and NRY variation using data from the Human Origins array showed that the modal NRY haplogroup in all Iranians and Parsis was J, with maximum frequency observed among the Parsis (freq=0.67; Figure 3a, Table S4). This is consistent with previous NRY haplogroup frequencies observed in Iranian Zoroastrian and non-Zoroastrian groups^68^. In particular, 8 of the 12 Iranian Zoroastrians from the city of Yazd belonged to NRY haplogroup J. In contrast, the modal NRY haplogroup among non-Zoroastrian Indian groups and groups in Pakistan was R (Figure 3a). In comparisons of NRY haplogroups among all Indian and Iranian groups, Parsis showed the lowest genetic distance with the Iranian Zoroastrian group in terms of F_ST_^40^ (F_ST_ = 0.026, p=0.157) and highest genetic distance with other Indian groups (Kharia and Tiwari; F_ST_ > 0.762, p < 0.0001) (Table S6).

**Figure 3.**
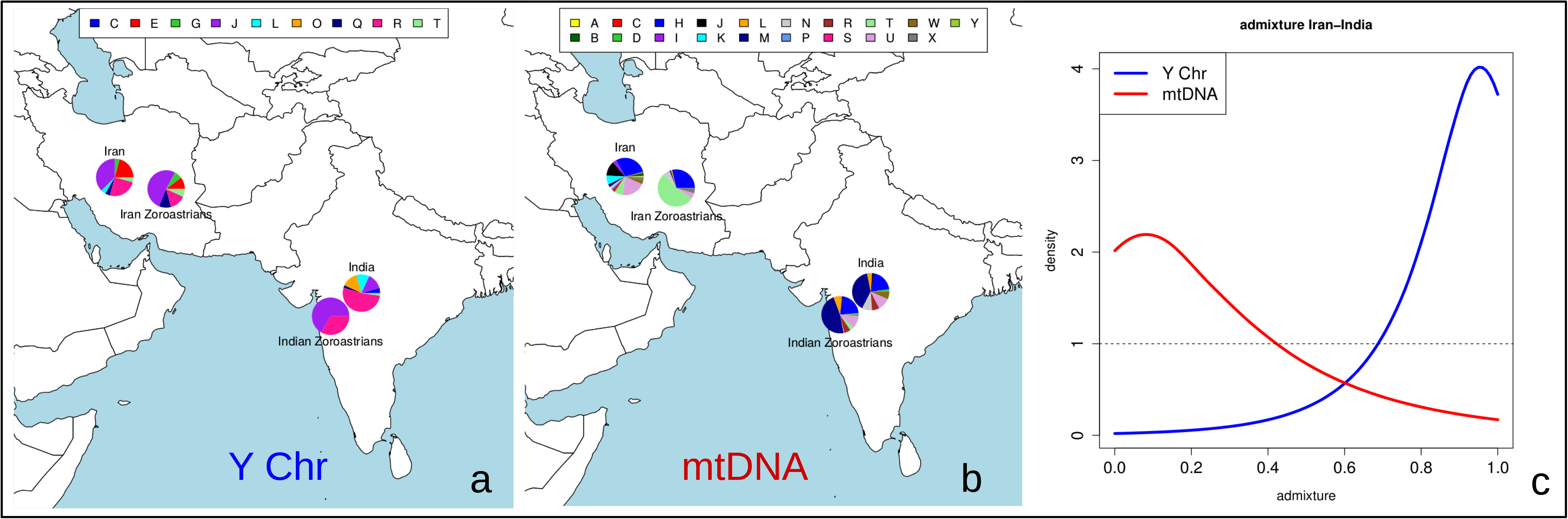
mtDNA and Y-chromosome variability in Iran and India. (a) NRY and (b) mtDNA macrohaplogroup frequencies in India, Parsis, Iran and Iran Zoroastrians using chip data in (a) and additional mitochondrial DNA sequencing in (b). (c) Posterior distribution of admixture proportions in lay Parsis assuming non-Zoroastrian Indian and Iranian lay Zoroastrian surrogate groups, using observed Mhg and Yhg values.

In contrast, the majority of individuals from India, Pakistan and the Parsis belonged to the same mtDNA haplogroup M, the modal mtDNA haplogroup in the Indian sub-continent (Table S5), also sharing the same modal sub-haplogroup M32’56 (Figure 3b). Parsis showed the highest genetic distance from the Iranian Zoroastrian group comparing mtDNA haplogroups (F_ST_ = 0.482, p < 0.0001), while having almost no genetic differentiation from other Indian groups (Kharia, Lodhi and Vishwabrahmin; F_ST_ < 0.0001, p > 0.487). The Pakistani groups were intermediary between groups from Iran and India, suggesting geographic continuity (Table S7). In Iran, however, since the batch of SNPs on the SNPchip typed are not representative of those diagnostic of haplogroups common in this region, most samples in the Zoroastrian and Iran groups are assigned to haplogroup H2a2a1 which is identical to that of the reference sequence (revised Cambridge reference sequence (rCRS) (Andrews et al., 1999).

Thus, to examine sex-biased admixture in Parsis in more detail we sequenced the mtDNA control region (positions 16,027 to 16,400^48^) in a larger sample of 79 Iranian lay Zoroastrians, 8 Iranian Zoroastrian priests, 121 lay Parsis, 71 Parsi priests, 46 non-Parsi Indians, and 193 non-Zoroastrian Iranians, and generated Y-chromosome haplotypes comprising 6 short tandem repeat (STR) and 12 biallelic loci^42,69^ in 76 Iranian lay Zoroastrians, 8 Iranian Zoroastrian priests, 122 lay Parsis, 71 Parsi priests, 41 non-Parsi Indians, and 172 non-Zoroastrian Iranians (Table S1). Using Y-chromosome binary polymorphism defined haplogroups (Yhg) and inferred mtDNA haplogroups (iMhg), these additional data showed that the Parsi priests sample has the lowest gene diversity values of all populations studied for both Y and mtDNA (Tables S12-S13), though we did not have enough data from Iranian Zoroastrian priests to make any analogous observation. Consistent with the Human Origins Y/mtDNA data, the iMhg and Yhg frequency-based pairwise F_ST_ values for these larger samples indicate that through the male line the lay Parsis have a closer relationship to the lay Iranian Zoroastrians, but through the female line they have a closer relationship to the non-Zoroastrians from India (Figure S10). However, no shared Y-chromosome STR+biallelic marker or mtDNA control region sequence haplotypes were shared between the Parsi priest and Iranian Zoroastrian priest samples, and all F_ST_ p-values and exact tests, whether based on Yhg, Y-haplotype, iMhg or mtDNA haplotype frequencies, indicated significant differentiation between these two.

Using the likelihood-based estimation of admixture (LEA) method of Chikhi et al.^55^ as implemented in the LEA software^70^ on Yhg and iMhg data, with the non-Zoroastrian Indians and Iranian lay Zoroastrians as surrogates for the two admixing source populations, we infer the most probable Iranian lay Zoroastrian contribution to the lay Parsis Y-chromosomes to be 96% (median = 86%, mean = 82%, 95% CI = 41% to 99%), whereas the most probable Iranian lay Zoroastrian contribution to Parsis mtDNA is 8% (Figure 3c; median = 26%, mean = 32%, 95% CI = 1% to 88%). More than ninety four percent of posterior estimates for Y-chromosome Iranian lay Zoroastrian contribution to the lay Parsis were higher than the posterior estimates for mtDNA Iranian lay Zoroastrian contribution to the lay Parsis in random samples drawn from each distribution. For comparison, the admixture proportion estimators mY, mC and mR for the Iranian lay Zoroastrian contribution to the lay Parsis^56,57,58^ gave very similar point estimates to the modal estimates obtained using LEA: For Yhg frequencies, mR = 0.94 (bootstrap SD = 0.093), mC = 0.93 (bootstrap SD = 0.11), mY = 0.96 (bootstrap SD =0.18). For iMhg frequencies, mR = 0.052 (Bootstrap SD = 0.15), mC = 0.12 (Bootstrap SD = 0.098), mY = 0.024 (Bootstrap SD = 0.16). For all 3 methods of admixture estimation the difference in estimated Y-chromosome and mtDNA contributions of the Iranian lay Zoroastrian contribution to the lay Parsis was highly significant.

### Inferring details of the Parsis priests

Our additional (i.e. non-Human Origins array) Y/mtDNA data defined 8 Y-chromosome haplogroups and 182 total Y-chromosome haplotypes when using biallelic and STR loci (Tables S12-S13) and 240 mtDNA haplotypes that clustered into 14 haplogroups using key HVS-1 mutations. These new data showed that the Parsi priests sample has the lowest gene diversity values of all populations studied in both Y and mtDNA, with the majority of the Parsi priest’s Y-chromosomes (86%) fall into either Yhg-1 or Yhg-28 (as defined in Figure S11). The distribution of STR-defined haplotypes within these haplogroups is characterized by the presence of a high frequency modal haplotype (>50%), with the remaining haplotypes being only a small number of mutation steps different from the modal haplotype (Figure S11). The exception to this is one ‘ outlier’ Yhg-28 chromosome that was found to be 9 mutation steps different from the nine-microsatellite defined Yhg-28 modal haplotype. These data are consistent with the majority of Parsi priests being patrilineal descendants of two male founders in the relatively recent past. Assuming that with the exception of the one Yhg-28 outlier, the modals are the ancestral haplotypes^61^ to all other chromosomes within each Yhg, we estimate the coalescence dates for Yhg-1 and Yhg-28 chromosomes are 37 generations (95% CI 19 to 61 generations) and 31 generations (95% CI 18 to 46 generations) respectively. Assuming a generation time of 28 years this translates to 1036 years (95% CI 532 to 1708 years) and 868 years (95% CI 504 to 1288 years) respectively. Noting that these two coalescence date estimates are not significantly different (only 63% of simulated dates for Yhg-1 are older than those for Yhg-28) we re-estimated the coalescence date assuming that both lineages originated at the same time by finding the mean ASD from the respective modal haplotypes for both clusters. This gave a combined coalescence date of 923 years (95% CI - 597 to 1277 years). When uncertainty in the mutation rate estimate is taken into account the 95% CI widens to 501 to 1782 years.

### Genetic regions showing evidence of selection in Zoroastrians relative to non-Zoroastrians

We calculated XP-EHH values for Iranian Zoroastrians and Parsis using other Iranians and Indians as reference populations (Figures S12-S13). Tables S14-S15 provide details for all the SNPs below and above quantiles 0.0001 and 0.9999 of the empirical distribution, respectively (see Methods), including the genes within those regions, or the flanking genes in the case of intergenic SNPs.

In the case of the Iranian Zoroastrians, most of the regions with the strongest signals of selection (positive XP-EHH values) are located in intergenic or intronic regions. Among these, some of the most significant SNPs (p<0.0001 based on a permutation procedure; see Methods) are located upstream from gene *SLC39A10* (Solute Carrier Family 39 Member 10) with an important role in humoral immunity^71^ or in *CALB2* (Calbindin 2), which plays a major role in the cerebellar physiology^72^.

With regards to the positive selection tests on Parsis versus India Hindu/Gujarati groups, the most significant SNPs were embedded in *WWOX* (WW Domain-Containing Oxidoreductase), associated with neurological disorders like epilepsy^73^, and in a region in chromosome 20 the *WFDC* (acidic protein WAP four-disulfide core domain) locus and other genes like *SPINT4*, *SNX21* or *TNNC2* (see Table S14 for a complete list). On the other hand, among the SNPs showing signatures of positive selection in the reference Indian population, two highly significant selection signals were identified: *LOC102467224* and *LOC283177*, with unknown functions.

## Discussion

Though recent studies have investigated the origins of different Jewish populations from India, like the Cochin Jews or the Bene Israel^74,75,76^, little is known about the genetic structure of the relatively isolated populations found mainly in India and Iran that practice Zoroastrianism, one of the oldest religions of the world. We present genome-scale genetic analyses of Zoroastrians from Iran and India, and provide genetic evidence for their historical exodus^3^.

Zoroastrians in both Iran and India are genetically differentiated from other groups in these countries, in Y-chromosome, mtDNA and autosomal patterns of variation (Figures 1,3, Figures S1-S5, S10, Tables S2-S3, S4-S7). For example, autosomal clustering using fineSTRUCTURE grouped all Parsis together with each other before merging with any other group, and merged 27 of 29 Iranian Zoroastrians with each other before merging them with any other group (Figure S1, Table S2). One of the remaining 2 Iranian Zoroastrians merged with 39 other individuals mainly from Lebanon and Turkey. The other merged with individuals we label as Iranian_B, which consists primarily of Bandari individuals, and shows a very similar genetic pattern and admixture history as this Iranian_B cluster (Figure S2, Table S16). Both of these two individuals were genetically distinct from the other Zoroastrians (Figure S2) suggesting these individuals were possibly mislabelled or recently converted to Zoroastrianism. The latter would suggest present-day Zoroastrians in Iran are not as closed a group today as previously reported^6^.

Excluding these two Iranian Zoroastrians, the remaining Zoroastrians in both Iran and India display a high level of genetic homogeneity; greater than any other Iranian and Indian group used in this study (Figure 1b). This is likely attributable to founder effects, bottlenecks and/or through some endogamy throughout the last millennium and up to the present day. These factors likely played a major role in the observed differences in autosomal DNA patterns between Iranian Zoroastrians and non-Zoroastrians from Iran, as analyses that attempt to mitigate these genetic isolation effects notably decrease the observed genetic differences between Iranian Zoroastrians and non-Zoroastrian Iranians (Figure 1d, Figure S4, Table S3). In contrast, our analyses to mitigate isolation effects do not drastically affect observed genetic differences between the Indian Zoroastrians (Parsis) and non-Zoroastrian groups from India, suggesting the different admixture histories of different Indian groups play a major role in shaping observed genetic differences among these Indian groups today (Figure 1c, Figure S5, Table S3).

In particular, we detect an admixture event in the Parsis dated to around 1030 CE (690-1390), between a source genetically similar to modern Indian groups and a second source best represented genetically by a ∼9,500 year old Neolithic farmer from Iran (Figure 2, Table S10). This Iranian source of introgression differs from the sources of admixture inferred in all other sampled Indian groups (Figure 2, Table S10). Our admixture date matches the historical records of a large-scale migration of Zoroastrians to India beginning in either 785 CE (Modi, 1905) or 936 CE^3^, providing genetic evidence for this period of migration and suggesting the migrants mixed with local females soon upon arrival. Our results suggest these migrations may have resulted in a single “pulse” of admixture occurring around 1030CE, though our dates are also consistent with multiple episodes of migration from around 690CE to 1390CE, which is difficult to disentangle given these sample sizes^27^. However, we only see evidence of Iranian origins in our Parsis and in no additional sampled non-Zoroastrian groups from India, which strongly suggests our admixture signal is due to the migration of Zoroastrians from Iran rather than related to historically documented trade routes between present-day Iran and India^5^ that would likely have included mixture among non-Zoroastrian groups.

That our approach inferred the Neolithic Iranian sample WC1 to be a better surrogate for the Iranian admixing source in the Parsis than any modern Iranian groups (including Iranian Zoroastrians) likely results from strong bottleneck effects and/or recent admixture events that have made modern Iranian groups look more genetically differentiated from the source group that migrated to India ∼17-44 generations ago. For example, when performing an alternative approach that attempts to mitigate genetic isolation effects within each modern Iranian and Indian group by disallowing genetic matching to members from the same assigned cluster (i.e. the “Non Indian/Iranian donors painting”; see Methods), this high aDNA contribution to Parsis is replaced by the modern Iranian Zoroastrians (Table S11, Figure S7). If we instead use the original approach that does not mitigate these isolation effects (i.e. the “all donors” painting in Figure 2, Table S10) but exclude WC1 as a surrogate, the highest contributing Iranian group to the Parsis is Iranian_A and not the Iranian Zoroastrians (Table S9). The fact that Iranian Zoroastrians are only favoured as the source of admixture in Parsis after mitigating isolation effects suggests that at least some of these effects in the Iranian Zoroastrians have occurred more recently than the migrations of Parsis to India ∼600-1300 years ago. In contrast, for the Parsis it is difficult to discern the extent to which their relative genetic homogeneity (e.g. Figure 1b) reflects recent isolation since admixture versus isolation effects occurring in their ancestry source from Persia prior to this admixture event.

Our mtDNA and NRY variation also shows clear evidence of contrasting maternal and paternal ancestry in Parsi individuals, consistent with previous studies which suggest that migration of the ancestors of the present-day Parsi population was largely sexually asymmetrical from Iran to India^77^. In particular, using Iranian lay Zoroastrians as a surrogate to this introgressing source in Parsis, the Iranian male contribution to the Parsis Y-chromosome gene pool with highest posterior probability is 96%, while the Iranian female contribution to the Parsis mtDNA gene pool with highest posterior probability is only ∼8% (Figure 3). Consistent with this, we infer the autosomal, sex-averaged contribution to be 61-76% using a variety of modern and ancient Iranian surrogate groups (Figure 2, Figure S7, Tables S9-S11). This supports Zoroastrianism being brought from Iran to India by a group of males, and/or that gene-flow into the Parsi community from the neighbouring Indian population was mainly female-mediated. Consistent with this, with the genetically estimated (see above) and historically attested arrival date of Parsis in India, and with the claim of patrilineal descent among Parsi priests, we infer that the majority of Parsi priests are descended from two male founders 923 years (95% CI - 597 to 1277 years) ago. This parallels the Jewish *kohanim* patrilineal priesthood, who claim descent from Moses’ brother Aaron, and display low Y-chromosome diversity; with most Y-chromosome STR haplotypes either belonging to or being only a small number of mutation steps away from a modal haplotype.

In Iranian Zoroastrians, we inferred a relatively old admixture event between sources best represented genetically by the Neolithic Iranian WC1 and modern-day Cypriots occurring in around 70 CE (range: 570 BCE-750 CE). While we infer admixture in each of our three other non-Jewish Iranian groups (Figure 2, Table S10), this admixture date in the Zoroastrians is significantly older, consistent with their long-standing isolation. The date uncertainty and ancient nature of this event prevents interpreting it in a clear historical context, but one intriguing possibility is that it might reflect mixture among groups joined via the allegiance of the Cypriots with Alexander the Great to help conquer the Persian Empire in 332 BCE. At any rate, interestingly our date range corresponds closely to that spanning the three major Persian empires (Achaemenid, Parthian, Sasanian) for which Zoroastrianism acted as official state religion (559 BCE-651 CE). Ancient DNA from these regions related to these ancient groups and others will greatly enhance our understanding of this older signal. Interestingly, when using only modern groups as surrogates and excluding WC1, GLOBETROTTER was not able to detect this older admixture event (Table S9). In this latter analysis, our model considered the Iranian Zoroastrians to be sufficiently genetically matched to a single modern group (Iranian_A) without requiring any other ancestry sources. Presumably this is because Iranian_A has similar genetic patterns to the Iranian Zoroastrians, with GLOBETROTTER inferring similar (but more recent) admixture 20-38 generations ago in Iranian_A between sources best represented by WC1 and modern-day Turkish groups. Our results here suggest that this similarity masks the older DNA contributions to the Zoroastrians. However, the combination of WC1 and other modern groups provides a better match to an ancestral source of the Iranian Zoroastrians than using only Iranian_A, enabling a clear signal of admixture (Tables S10-S11, Figure 2, Figure S7). This reveals how adding even small numbers of ancient samples, particularly those less affected by recent admixture, can increase power and insights in population genetic history inference, even if those ancient samples are substantially older than the time period under study, as is the case here with WC1 living over 7,000 years earlier.

Genetic isolation and endogamous practices can be associated with higher frequencies of disease prevalence. For example, there are reports claiming a high recurrence of diseases such as diabetes^78^ among the Iranian Zoroastrians, and Parkinson’s^79^, colon cancer^80^ or the deficiency of G6PD^81^, an enzyme that triggers the sudden reduction of red blood cells, among the Parsis. Researchers have argued that in addition to these demographic effects, selection can also play a role in the increase of rare disorders or other phenotypes, as has been previously reported for the Ashkenazi Jews^82,83^. Therefore, identifying regions under positive selection in the Zoroastrian populations may be helpful to understand the prevalence of diseases or distinct phenotypic traits in the community. Supporting this, using XP-EHH^33^ comparing Zoroastrians to non-Zoroastrians, we have identified some regions that might have been under selection specifically in the Zoroastrians (p<0.0001 based on a permutation procedure; see Methods), as well as putative selection in the non-Zoroastrian reference groups. Some of these regions contain genes that have been associated with different diseases, including cancers, like *DEC1* associated with esophageal cancer^84^ and positively selected in Iranian non-Zoroastrians, or *WWOX*, associated with spinocerebellar ataxia^85^ and epilepsy^73^ and positively selected in Indian Zoroastrians. However, a permutation study that re-assigned Zoroastrians and non-Zoroastrians randomly to two groups and then tested for selection between these groups gave very similar magnitudes of XP-EHH scores to that seen in our non-permuted data (Figure S13), warranting caution in interpreting these findings. A larger cohort would be needed to corroborate their significance, coupled with exhaustive epidemiological studies. Nonetheless, they represent a first insight into understanding genetic predisposition and/or resistance to disease in these groups that could form the basis for targeted medical approaches in these isolated groups.

In summary, in this work we explore the genetic landscape and structure of India and Iran and provide genome-wide genetic evidence that the Parsis descend from an admixture event between ancestral groups consisting predominantly of males with Iranian-related ancestry and females with Indian-related ancestry. For the first time, we date this event in ancestral Parsis to around 1030 CE, in agreement with historical records, and also provide new evidence of a much older admixture event in Iranian Zoroastrians dated to around 74 CE with an unknown historical explanation. We also demonstrate that Zoroastrians in both countries are genetically homogeneous populations differentiated from other population living locally; likely in part due to strict religious rules that discourage intermixing with non-Zoroastrians. Further work is required to help understand whether the genetic differences attributable to this isolation correlate with observed differences in disease phenotypes between these communities and other local groups.

## Supplemental Data

Supplemental Data include 13 figures and 16 tables.

## Acknowledgements

SL is supported by BBSRC (Grant Number BB/L009382/1). GH is supported by a Sir Henry Dale Fellowship jointly funded by the Wellcome Trust and the Royal Society (Grant Number 098386/Z/12/Z) and supported by the National Institute for Health Research University College London Hospitals Biomedical Research Centre. LvD is supported by CoMPLEX via EPSRC (Grant Number EP/F500351/1). We are grateful to Zoroastrian Studies (Bombay) and in particular to Mr Khojeste P. Mistree, for the facilitation of the project in India and the introduction to priests in all the locations where samples were obtained. We also thank Mrs Firoza Punthakey Mistree, Ms Tashan Mistree and the late Mrs Shehnaz Neville Munshi for her involvement in the samples collection in India and Iran. Work on the genetics of these samples is covered by UK ethics committee London Bentham REC (formally the Joint UCL/UCLH Committees on the Ethics of Human Research: Committee A and Alpha), REC reference number 99/0196, Chief Investigator = Mark Thomas. We also thank the Children’s Hospital of Philadelphia for genotyping the samples on the Human Origins array.

